# Identification of misclassified ClinVar variants using disease population prevalence

**DOI:** 10.1101/075416

**Authors:** Naisha Shah, Ying-Chen Claire Hou, Hung-Chun Yu, Rachana Sainger, Eric Dec, Brad Perkins, C. Thomas Caskey, J. Craig Venter, Amalio Telenti

## Abstract

There is a significant interest in the standardized classification of human genetic variants. The availability of new large datasets generated through genome sequencing initiatives provides a ground for the computational evaluation of the supporting evidence. We used whole genome sequence data from 8,102 unrelated individuals to analyze the adequacy of estimated rates of disease on the basis of genetic risk and the expected population prevalence of the disease. Analyses included the ACMG recommended 56 gene-condition sets for incidental findings and 631 genes associated with 348 OrphaNet conditions. A total of 21,004 variants were used to identify patterns of inflation (i.e. excess genetic risk). Inflation, i.e., misclassification, increases as the level of evidence in ClinVar supporting the pathogenic nature of the variant decreases. The burden of rare variants was a main contributing factor of the observed inflation indicating misclassified benign private mutations. We also analyzed the dynamics of re-classification of variant pathogenicity in ClinVar over time. The study strongly suggests that ClinVar includes a significant proportion of wrongly ascertained variants, and underscores the critical role of ClinVar to contrast claims, and foster validation across submitters.

## INTRODUCTION

Currently, more than 60,000 clinical genetic tests are offered from more than 1600 clinics and laboratories (https://www.genetests.org/). While genetic testing is a powerful diagnostic tool, there are several challenges for interpretation and reporting of findings. Some of these challenges include the accuracy of variant calling, identification of pathogenic variants and interpretation of low-penetrant variants. Different laboratories have developed different protocols to handle these challenges, which leads to inconsistencies in the classification of variants and a bias toward overestimating pathogenicity^1^.

For many variants, the assignment of clinical significance reflects historical guidelines and evidence available at the time of the original interpretation. However, as additional information becomes available, the interpretation of pathogenicity of genetic variants may change. Data-sharing efforts have shown that 17% of the variants with clinical interpretations submitted by more than one laboratory had conflicting interpretations^2^. The American College of Medical Genetic and Genomics (ACMG) and the Association for Molecular Pathology (AMP) issued guidelines to support a standardized approach to variant classification^3^. Initiatives to curate existing knowledge and improve variant interpretation have been put in place. The NIH-based partnership between ClinVar and ClinGen is an example of such an initiative ^2,4^. ClinVar implements a ranking system to denote the quality associated with each submission to the database. For example, a three- or four-star submission comes from “expert panel” and “practice guidelines” submitters, which are the most ClinGen-trusted sources for variant interpretation. Challenges lie ahead, as even the implementation of ACMG-AMP guidelines led to only a 34% concordance on the reporting of 99 variants across laboratories. After consensus discussions and detailed review of the ACMG-AMP criteria, concordance increased to 71% ^1^.

Leveraging knowledge from shared data by categorizing variants based on clinical significance, the number of submitters, and their assertion criteria is an important step towards accurate interpretation of variant pathogenicity and diagnosis. However, there is a need for additional methods to detect misclassified pathogenic variants. Also, it is crucial to identify variants with low clinical penetrance (i.e. proportion of individuals with a variant that develop the disease or clinical symptoms) for proper clinical reporting. For example, variant C282Y in the *HFE* gene associated with hereditary hemochromatosis was thought to be the main pathogenic variant^5^; however, as more individuals were genotyped, the variant’s high population frequency appeared more compatible with low penetrance^6^.

Here, we revisit the topic of assignment of pathogenicity to a variant by assessing expected disease prevalence and observed genetic risk (genetic risk is defined here as the number of individuals that are at disease risk based on pathogenic variants and mode of inheritance) in a population. We used recent data from whole genome sequencing of 8,102 unrelated subjects to identify individuals with clinically significant variants from ClinVar. We identify disease conditions with inflated values, i.e. the cumulative frequency of the disease risk variants in the population far exceeds the expected prevalence of disease. If we assume that the variants are truly pathogenic and have complete penetrance, then the frequency of diagnosed individuals should match the estimated population prevalence of the diseases. If the genetic basis and etiology of a disease is not well understood (incomplete knowledge on associated genes, variants and other factors), the currently known variants will only explain some of the disease prevalence, i.e., they will appear deflated. We performed the disease prevalence analysis as a function of pathogenic variants listed in ClinVar separately for the well-curated 56 genes and associated conditions recommended by the ACMG for reporting of secondary findings in clinical genome-scale sequencing^7,8^ (herein referred to as ACMG-56), and for genes with available population prevalence information reported in Orphanet/OrphaData, a data source on rare diseases^9^.

## MATERIALS AND METHODS

In the study, we used whole genome sequences from 8,102 unrelated individuals sequenced at a mean coverage of 30X. The samples are described in detail by Telenti *et al.*^10^: participants were representative of major human populations and ancestries and the study population was not ascertained for a specific health status. To avoid analyzing potentially inaccurate variant calls that lie within the areas of the genome prone to sequencing errors^11^, the analysis was focused on variants that fell within the high confidence sequencing regions of the genome^10^,^12^,^13^. For the disease prevalence analysis, we calculated frequencies of individuals at genetic risk using variants deposited in ClinVar^14^. We performed disease prevalence analysis for two groups of conditions: 1) a well-curated list of the recommended ACMG 56 gene and associated conditions to report for incidental findings (referred to here as “ACMG-56”) ^7^, and 2) a list of rare conditions collected by a consortium in a reference portal called Orphanet/Orphadata (http://www.orphadata.org/, http://www.orpha.net/). To compare observed genetic risk and expected population prevalence, we used only the conditions with two or more at-risk individuals observed in the study. We calculated fold-change for genetic-risk compared to population prevalence per condition using the formula: observed/expected.

## ACMG-56 Conditions

For each of the ACMG-56 conditions, we searched OrphaNet, GeneReview and other published sources for the available population prevalence of disease. In case of multiple prevalence information available, we chose the maximum prevalence for the purpose of the study.

## OrphaNet Conditions

We selected Orphanet conditions that had at least one associated ClinVar variant and had a defined mode of inheritance and population prevalence information available (i.e. prevalence type of "Point prevalence" and "Prevalence at birth"). Only the conditions that had the following mode of inheritances were considered: autosomal dominant, autosomal recessive, X-linked recessive, and X-linked dominant. In case of multiple population prevalence information available, we chose the maximum prevalence.

## ClinVar Variants

We used the ClinVar XML file (version: ClinVarFullRelease_2016-05.xml.gz) to obtain the disease-associated variants. We filtered out variants that were tagged as “haplotype”, “compound heterozygotes” and “phase unknown”. Following the ACMG guidelines for clinical interpretation, we removed variants with greater than 5% allele frequency. If a variant was called both protective and pathogenic, we did not include it in the analysis. We divided variants deposited in ClinVar using ClinVar/ClinGen’s ranking system2 into four sets: *Set-1* included pathogenic and likely pathogenic (P/LP) variants with ClinVar star 2+ (i.e. multiple submitters with assertion criteria, expert panel or practice guideline). *Set-2* included P/LP variants with ClinVar star 1 (i.e. one submitter with assertion criteria). *Set-3* included P/LP variants with ClinVar star 0 (i.e. submitter without assertion criteria). *Set-4* included variants with conflicting clinical significance. Each of the sets of variants was used separately to perform the disease prevalence analysis for ACMG-56 and Orphanet conditions.

## Disease-specific Allele Frequency Threshold

To identify and remove potentially benign variants from the sets above, we applied disease-specific allele frequency (dAF) threshold per condition. We defined the dAF threshold as: 1) For autosomal/X-linked dominant conditions, assuming there is one highly penetrant variant causing 100% of the disease cases, then the frequency of heterozygous should not be greater than the disease prevalence. Thus, the AF for the variant should not exceed ½*(disease prevalence). 2) For autosomal/X-linked recessive conditions, assuming there is one highly penetrant variant causing 100% of the disease cases, then the frequency of homozygous recessive should not be greater than the disease prevalence. Thus, the AF for the variant should not exceed the square root of the disease prevalence.

To account for penetrance, these formulas can be generalized to: 1) For dominant conditions, dAF threshold = ½*(disease prevalence)*(1/penetrance). 2) For recessive conditions, dAF threshold = sqrt(disease prevalence*(1/penetrance)).

However, for the study, since one of our goals is to highlight conditions with inflated genetic risk, we use a more stringent threshold assuming 100% penetrance. We applied the threshold to variants observed in more than one individual in the study.

## Change in ClinVar variant classification

To investigate how ClinVar variant classification changed over time, we compared the version used in our analysis, May 2016, to August 2015 version. We selected variants that were common between the two versions and that had clinical significance terms recommended by ACMG (i.e. pathogenic, likely pathogenic, benign, likely benign, and VUS). We grouped together variants that had a clinical significance of pathogenic and/or likely pathogenic as P/LP. Similarly, we grouped together variants with clinical significance of benign and/or likely benign as B/LB.

## RESULTS

We performed genetic screening of 8,102 unrelated individuals, whose whole genomes were sequenced with a mean coverage of 30X ^10^. For genetic screening, we used disease-associated variants deposited in ClinVar^2^. We divided the variants into four sets based on ClinGen’s ranking system using clinical significance and review stars (see Methods for more details). *Set-1* includes 1,879 pathogenic and likely pathogenic (P/LP) variants with ClinVar star 2+ (i.e. multiple submitters with assertion criteria, expert panel or practice guideline). *Set-2* includes 14,975 P/LP variants with ClinVar star 1 (i.e. one submitter with assertion criteria). *Set-3* includes 24,759 P/LP variants with ClinVar star 0 (i.e. submitter without assertion criteria). *Set-4* includes 5,735 variants with conflicting clinical significance. Below we present the analyses for ACMG-56 and for Orphanet genes and conditions.

## ACMG-56

Twenty-four sets of medical conditions and 56 genes are represented in ACMG-56. There are 11,084 ClinVar variants associated with the ACMG-56 genes (n_set-1_ = 1333, n_set-2_ = 4421, n_set-3_ = 3670, and n_set-4_ = 1660). It should be noted that 12 of the 24 sets of medical conditions did not have any set-1 variants, which are the most reliable according to the ClinGen/ClinVar ranking. Lynch syndrome was the only condition that had substantially more P/LP variants in set-1 compared to P/LP variants in set-2 and set-3.

In the study population, we observed 21,465 variants with allele frequency < 5% in the coding regions of the 56 ACMG genes, including 5,818 missense and 88 loss-of-function (LoF) variants. Of these, 1,134 variants were matched to ClinVar records: n_set-1_ = 23, n_set-2_ = 162, n_set-3_ = 51, and n_set-4_ = 898. Screening for the P/LP variants from set-1, set-2 and set-3, we observed that 3.95% of the individuals would be predicted at risk for disease (herein referred to as “genetic risk”) for 18 of the 24 ACMG-56 conditions, and 5.34% of the individuals were carriers for 19 of the 24 ACMG-56 conditions. This is within the estimated range (1.5-6.5%) of screened individuals that would have an incidental finding for the ACMG-56 ^15^. Five individuals (0.06%) in the study were at genetic risk for two ACMG-56 conditions.

We wanted to investigate if the variant ranking (here, broken down by variant sets) is indicative of misclassified variants. Thus, using variants from each set separately, we compared the observed genetic risk to the reported population prevalence for the conditions (Figure 1A). We would expect by using subsets of variants, the observed genetic risk would be lower-bound if the variants were truly pathogenic and with high penetrance. While there was no inflation (heuristically defined as more than 10-fold increase; see Methods) in the observed genetic risk using variants from set-1, we observed inflated genetic risk for several conditions using variants from set-2 and set-3. This may indicate that some of the variants have either low penetrance or inaccurate pathogenicity assignment. As we sequentially added ranked sets of variants to calculate genetic risk, the fold-change of observed genetic risk compared to population prevalence gradually increases (Figure 1B). This suggests that with the addition of more variants with lower ranks, more misclassified variants and/or variants with low penetrance accumulate and contribute to the inflation. We found 6 conditions with more than 10-fold increase (i.e. inflated) when using P/LP variants from set-1, set-2 and set-3 (two of the same conditions were inflated in both set-2 and set-3; Figure 1A). These conditions included malignant hyperthermia susceptibility, multiple endocrine neoplasia type 1, Romano-ward long QT and Brugada syndromes, Ehlers-Danlos syndrome (vascular type), Retinoblastoma, and hereditary paraganglioma-pheochromocytoma syndrome. Below we discuss several of the conditions.

**Figure 1:**
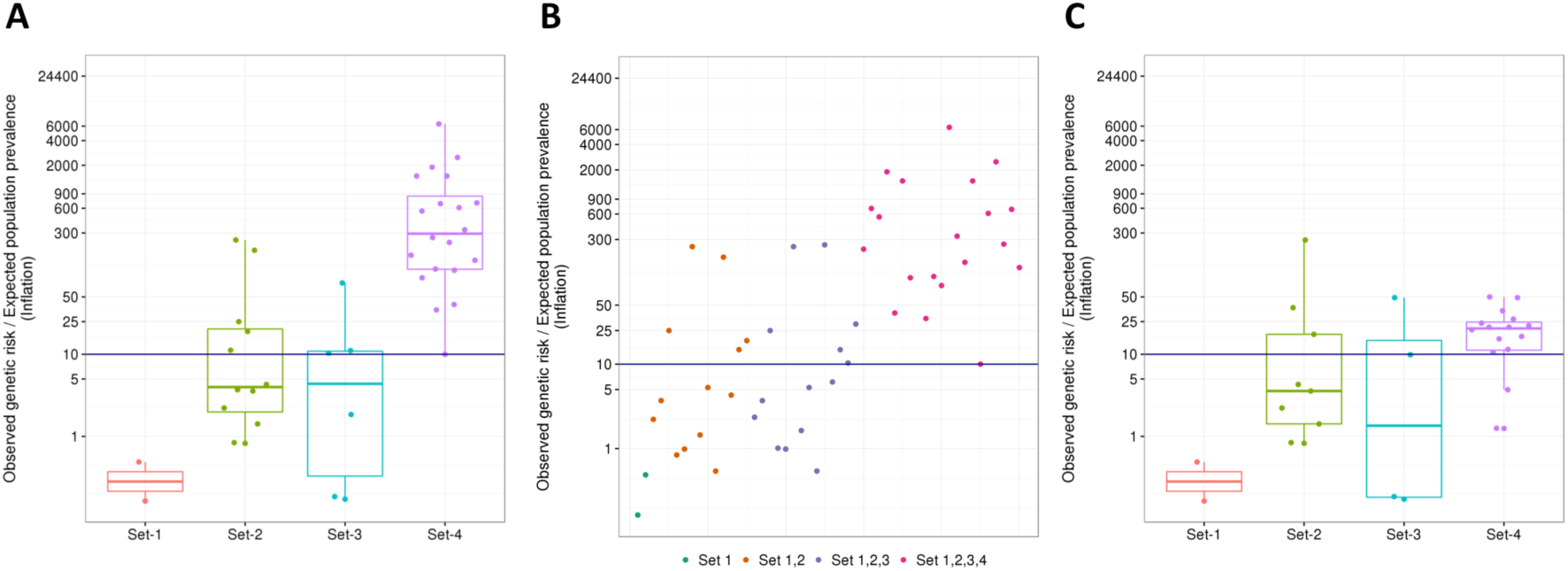
Genetic risk in ACMG-56 conditions. Fold-change of observed genetic risk over expected population prevalence using ClinVar variant sets for the ACMG-56 conditions. Each point represents a condition; each condition may be represented in more than one set. The navy-blue line at a fold-change of 10 (i.e. inflation) indicates a theoretical penetrance of 10%. Observations above this line are highly suggestive of misclassified variants. A) Fold-change was calculated using variants per variant set: Set-1 consists of variants with 2 or more ClinVar review stars (i.e. two or more submitters with assertion criteria, expert panel and practice guideline); Set-2 consists of variants with 1 star (i.e. one submitter with assertion criteria); Set-3 consists of variants with 0 star (i.e. submitter with no assertion criteria submitted in ClinVar); Set-4 consists of variants with conflicting interpretations of pathogenicity. B) Fold-change was calculated by using variants cumulatively from each set; i.e., Set-2 includes Set-1 variants, set three includes Set-1 and 2 variants, Set-4 includes all variants. C) Fold-change was re-calculated after variants were filtered for disease-specific allele-frequency thresholds.

With variants of conflicting interpretations from set-4, the observed genetic risk inflated massively (Figure 1A). This is a strong indication of misclassified variants in the set of conflicting interpretation, in particular as inflation goes far beyond what could be assumed to reflect low penetrance. There may be a few exceptional variants within the set that may be pathogenic; however, without supporting data they should be removed from consideration.

A recommended criterion to identify benign variants beyond ACMG-AMP’s criteria of greater than 5% allele frequency (AF) threshold is to develop disease-specific allele-frequency (herein referred to as dAF) thresholds^1^. After filtering out variants using dAF, we observed an overall decrease in inflation, especially using set-4 variants (Figure 1C), thus, confirming that a large proportion of the set-4 variants are benign. However, the genetic risk of three conditions (Romano-ward long QT and Brugada syndromes, hereditary Paraganglioma-Pheochromocytoma syndrome, and malignant hyperthermia susceptibility) was still inflated using P/LP variants. Individually, variants for these disorders are rare; however, collectively, they add up to show an increase in the observed genetic risk compared to the estimated population prevalence. These conditions with inflated risks are discussed in detail below, and as a model case we also describe a well-studied condition – hereditary breast and ovarian cancer.

Overall, of the 1134 ClinVar variants from all sets (236 in set-1,2,3) observed in 8102 individuals, we removed 305 (27%) variants from all sets (13, 5.5% in set-1,2,3) using the dAF filter. Of the 24 ACMG conditions, the genetic risk of 6 were inflated before the dAF filter (set-1,2,3 only) and 3 were still inflated after the filter. Therefore, for the critical sets (sets 1, 2, 3) only few variants appear responsible for the inflation.

### Hereditary Breast and Ovarian Cancer (HBOC) as a model

To test that the disease estimates for well-studied conditions are as expected, we studied the frequency of individuals at genetic-risk for hereditary breast and ovarian cancer (HBOC). HBOC is estimated to have a disease prevalence of 0.2%-0.3% in the general population^16^. Variant classification for *BRCA1/2* showed high concordance across seven established clinical testing laboratories http://meeting.ascopubs.org/cgi/content/abstract/34/15_suppl/1592). Using P/LP variants, we observed 24 individuals with 23 variants to be at genetic risk for HBOC (0.3% of the individuals); i.e. the frequency is as expected. As of September 2016, all of the 23 variants are classified as either 2 or 3 stars in ClinVar (n=8 and n=15 respectively).

### Romano-Ward Long QT Syndrome and Brugada Syndrome

These syndromes are two phenotypes of the sodium channel disease that are associated with abnormal QT interval and sudden cardiac death. These conditions are inherited as autosomal dominant disorders. We found the observed genetic risk for these syndromes inflated several fold compared to expected population prevalence of the disease (1358 versus 50 per 100,000 using dAF filtered P/LP variants from set-1, set-2 and set-3). In the study, we observed 75 variants in 110 individuals for the three main disease-associated genes (*KCNH2*, *KCNQ1* and *SCN5A*). These genes account for 70% to 75% of congenital LQTS cases^17^. The inflation suggests that many reported variants are of very low penetrance or represent benign private mutations. This observation is in line with the recent report of Van Driest *et al.*^18^ describing the low concordance in designating *SCN5A* and *KCNH2* variants as pathogenic across three laboratories and the ClinVar database. Strict application of ACMG criteria would have preempted this excess ^19^. On the other hand, LQTS may be more common than the estimated prevalence, since many individuals remain asymptomatic ^20,21^.

### Hereditary Pheochromocytoma-Paraganglioma

This is a rare condition characterized by growth of benign tumors in paraganglia, and with an inheritance pattern of autosomal dominant. We observed 6 individuals in the study carrying 6 different variants (all from set-2) in the disease-associated *SDHB* and *SDHD* genes (observed genetic risk of 74 in 100,000 individuals). The population disease prevalence is not precisely known but the rate of incidence is approximately 0.3 in 100,000 individuals^22^. Disease penetrance for both the genes has the highest score (>40% chance) and clinical severity of 2 (i.e. reasonable possibility of death-or major morbidity) according to ClinGen guidelines for actionability assessment for ACMG-56 4. According to Benn et al.^23^, by the age of 40 years the penetrance for *SDHD* gene is 68%. We found that four of the six variants lack sufficient information in ClinVar to determine pathogenicity. One variant (rs570278423) in *SDHB* was previously reported to be associated with paraganglioma syndrome^24^; however, the ClinVar submitter (ClinVar ID: RCV000183223) notes that the possibility of the variant being a rare benign mutation cannot be excluded. Reporting such variants that are not well recognized and do not have enough information available to clarify the association could lead to false reporting.

### Malignant hyperthermia susceptibility

This is a disorder of skeletal muscle calcium regulation and is inherited in an autosomal dominant manner. This disorder is triggered by reactions to anesthetics and thus the exact population prevalence is unknown. However, prevalence in individuals undergoing surgery in a New York hospital was estimated to be 1 in 100,000 for adults and 3 in 100,000 in children^25^. Incidence range is estimated much higher to be 1 in 30,000 to 50,000 uses of anesthetics^25^. Using P/LP variants in *CACNA1S* and *RYR1* genes from set-1, set-2, and set-3 (after applying dAF threshold), the observed genetic risk was inflated to 86 in 100,000 individuals. A total of 7 P/LP variants in only the *RYR1* gene were observed in 7 individuals.

*RYR1* is also related to several distinct myopathies, including central core disease, multiminicore disease, congenital fiber type disproportion, centronuclear myopathy, King-Denborough Syndrome, and nemaline myopathy^25^. Some of the myopathies are inherited in an autosomal dominant manner and some are inherited in autosomal recessive manner^26^. The confluence of several disorders, two modes of inheritance, and a disorder triggered by an exogenous exposure (anesthetics) leads to great imprecision in estimates of population prevalence and genetic risk.

## OrphaNet Conditions

Next, we performed a similar prevalence analysis on OrphaNet conditions with at least one variant in any of the four sets, a stated mode of inheritance and disease prevalence information. There are 14,400 ClinVar variants (n_set-1_ = 504, n_set-2_ = 5006, n_set-3_ = 7852, and n_set-4_ = 1038) in 631 genes associated with 348 OrphaNet conditions.

In the study population, we observed 179,426 variants with allele frequency < 5% in the coding regions of the 631 genes, including 45,811 missense and 1,057 LoF variants. Of these, 1,682 variants were matched to ClinVar records: n_set-1_ = 109, n_set-2_ = 552, n_set-3_ =466, and n_set-4_ =555. Screening for the P/LP variants from set-1, set-2 and set-3, we observed 14.1% of the individuals with genetic risk for at least one of the 67 OrphaNet conditions, and 64.5% of the individuals were carriers for at least one of 220 OrphaNet conditions. Lazarin *et al.* identified 24% individuals of a large, ethnically diverse population as carriers for 108 Mendelian disorders^27^. While the present study is based on whole genome sequencing and not limited to predefined sets of variants and disorders, the massive increase in carriers may alert of a significant number of misclassified variants.

As with the ACMG-56 conditions, we compared the observed genetic risk to the reported population prevalence for the Orphanet conditions separately for each set (Figure 2A). We observed one condition, hereditary chronic pancreatitis (HCP), with more than 10-fold higher observed genetic risk compared to the expected population prevalence using variants from set-1. These P/LP variants (n=3) are in *CFTR*, which is known to be associated with the autosomal dominant HCP and the autosomal recessive cystic fibrosis. The mode of inheritance for the *CFTR* gene and its associated condition HCP is incorrectly assigned as autosomal dominant in Orphanet. Given this discrepancy, the observed genetic risk was inflated. Using set-2 and set-3 variants, we observed 14 conditions in each set with inflated genetic risk. Using P/LP variants from the union of set-1, set-2 and set-3, we found 24 conditions with inflated observed genetic risk. As observed with the ACMG-56 conditions, genetic risk using set-4 variants inflated massively for 50 Orphanet conditions; strongly suggesting the inclusion of misclassified variants (Figure 2B).

**Figure 2:**
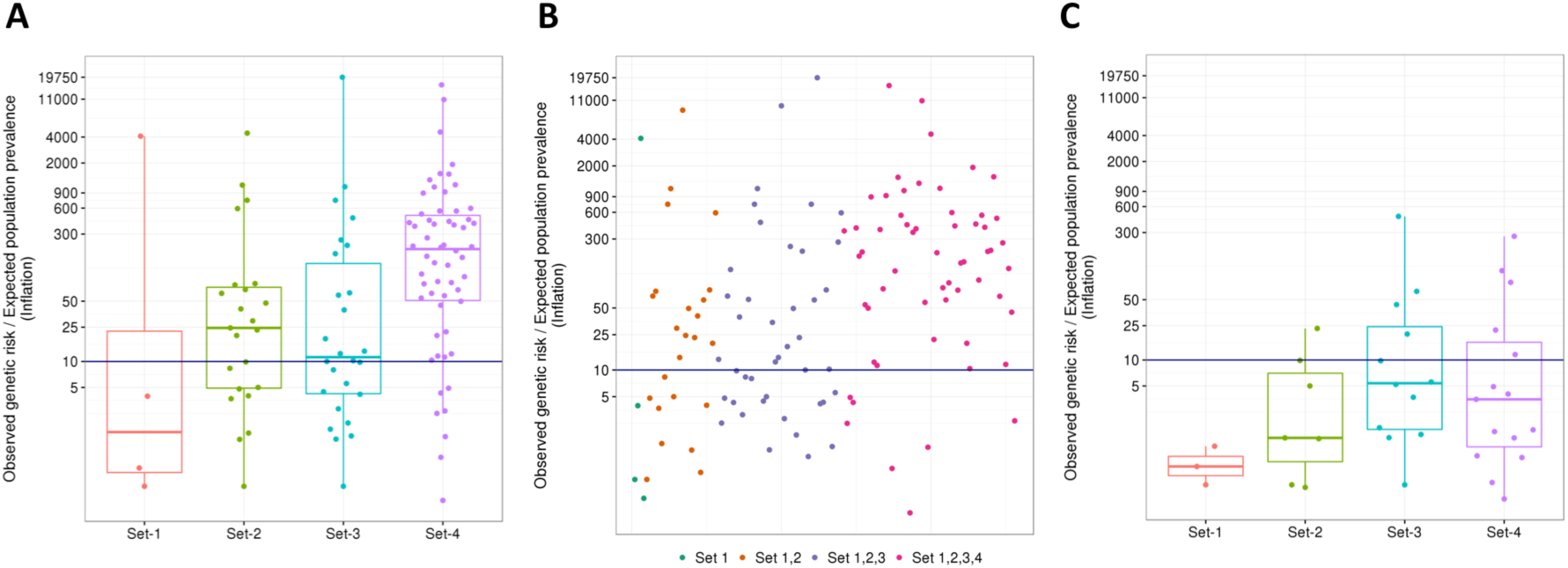
Genetic risk in Orphanet conditions. Fold-change of observed genetic risk over expected population prevalence using variant sets from ClinVar for the Orphanet conditions. Each point represents a conditon; each condition may be represented in more than one set. The navy-blue line at a fold-change of 10 (i.e. inflation) indicates a theoretical penetrance of 10%. Observations above this line are highly suggestive of misclassified variants. A) Fold-change was calculated using variants per variant set: Set-1 consists of variants with 2 or more ClinVar review stars (i.e. two or more submitters with assertion criteria, expert panel and practice guideline); Set-2 consists of variants with 1 star (i.e. one submitter with assertion criteria); Set-3 consists of variants with 0 star (i.e. submitter with no assertion criteria submitted in ClinVar); Set-4 consists of variants with conflicting interpretations of pathogenicity. B) Fold-change was calculated by using variants cumulatively from each set; i.e., Set-2 includes Set-1 variants, set three includes Set-1 and 2 variants, Set-4 includes all variants. C) Fold-change was re-calculated after variants were filtered for disease-specific allele-frequency thresholds.

We used dAF threshold to filter out additional potentially benign and low penetrant variants from the variant sets (see Methods). We observed an overall decrease in inflation (Figure 2C). Applying the filter, we not only removed potentially benign variants but also filtered out variants due to inaccurate disease information. For example, the relatively common variant rs6025 (also known as R506Q mutation; MAF = 0.02) is incorrectly associated with factor V deficiency (prevalence of 1 in 1,000,000) ^28^. The mutation is well known for its association with a more common clotting disorder called factor V Leiden, not to be confused with factor V deficiency, a bleeding disorder. Another example is the successful removal of variants in the *CFTR* gene that were associated with HCP.

Even though the dAF filter removed most of the noise, there were five autosomal/X-linked dominant conditions and one X-linked recessive condition with inflated genetic-risk using P/LP variants collectively from set-1, set-2 and set-3. The conditions were: central core disease, Cushing syndrome due to macronodular adrenal hyperplasia, hereditary breast and ovarian cancer syndrome (note: in addition to its associated genes *BRCA1* and *BRCA2* in ACMG-56, ClinVar contains associated variants in the *RAD51D*. Also, the prevalence information is an older estimate in Orphanet), hereditary chronic pancreatitis, X-linked Alport syndrome, and Menkes disease.

Overall, of the 1,682 ClinVar variants from all sets (1127 in set-1,2,3) observed in 8102 individuals, we removed 348 (21%) variants from all sets (69, 6% in set-1,2,3) using the dAF filter. Of the 348 Orphanet conditions, the genetic risk of 24 were inflated before the dAF filter (set-1,2,3 only) and 6 were still inflated after the filter. As it was the case for ACMG-56 conditions, only few variants appear responsible for the inflation in sets-1,2,3.

## Changes of classification of ClinVar variants over time

The classification of variants in ClinVar evolves over time as a reflect of better understanding of pathogenicity, but also as a result of more population representative frequency data emerging through large sequencing efforts. To better understand the evolution of the assignment of pathogenicity classification to variants, we identified the directionality of the changes, i.e., whether a pathogenic/likely pathogenic (P/LP) variant was reclassified as a variant of uncertain significance (VUS), benign/likely benign (B/LB) or of conflicting pathogenicity. For this purpose, we compared the August 2015 release to the May 2016 ClinVar release (see Methods). We observed that a majority of the variants (n=81,823 of 83,534 variants; 97.95%) did not change its clinical significance classification over the period of 9 months. For the 1,711 (2.05%) variants that were re-classified, we noted the directions of change (Figure 3).

**Figure 3:**
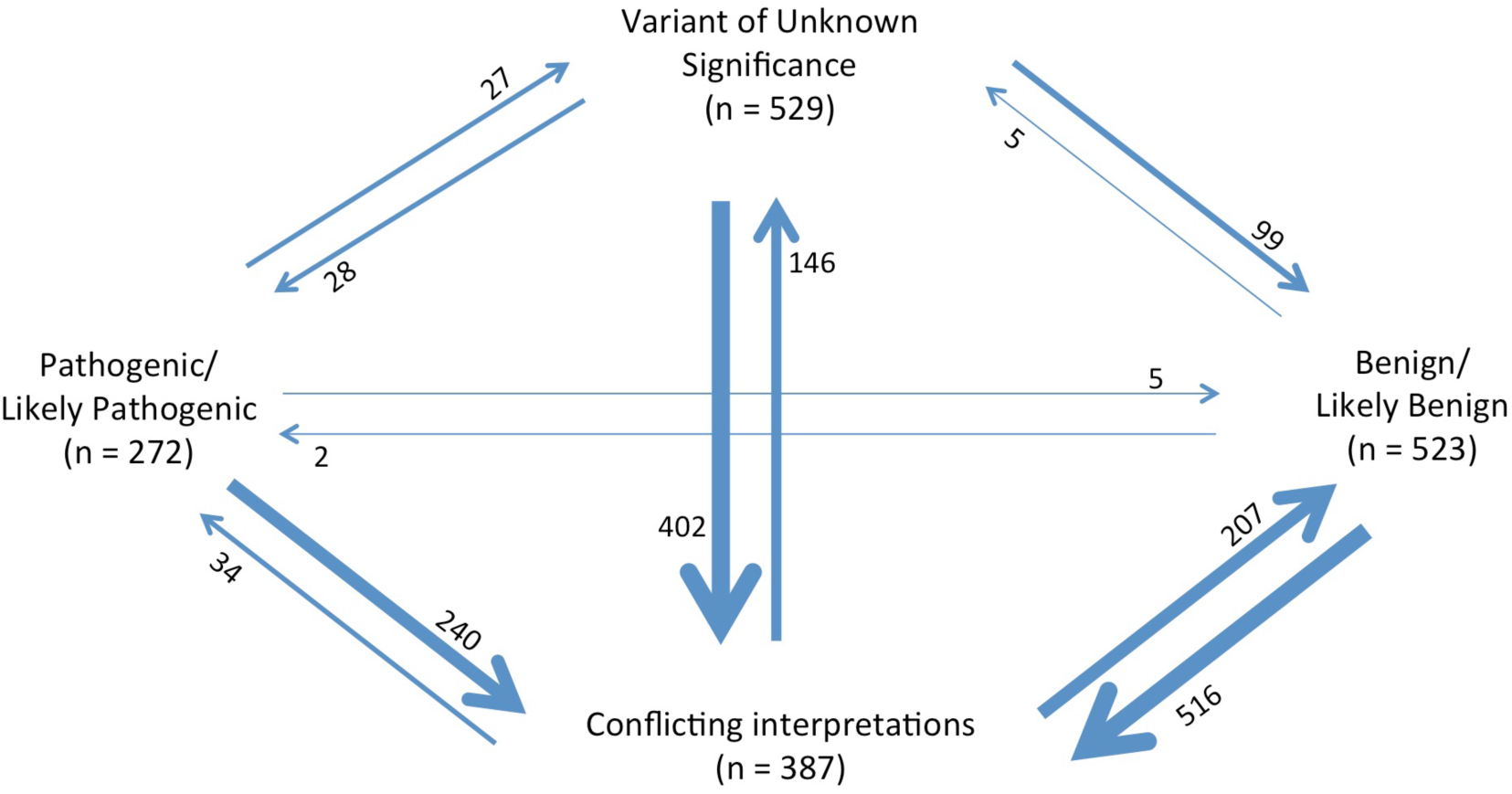
Change in ClinVar variant classification from August 2015 to May 2016. In the study period, 1,711 ClinVar variants changed classification. Predominantly, variants were reclassified as “conflicting interpretation” (n=1158; 67.68%). Only 64 variants (3.74%) were reclassified as pathogenic or likely pathogenic. Thickness of the arrows corresponds to the number of variants reclassified.

The most dynamic classification was “conflicting interpretations of pathogenicity” with 387 of 4847 “conflicting” variants (outbound, 7.98%) changing the interpretation to other classifications, while 1158 of 1711 reclassified variants (inbound, 67.68%) were reclassified as “conflicting interpretations of pathogenicity”.

In contrast, the P/LP classification was the least dynamic set in terms of reclassified variants with only 272 of 35,462 (outbound, 0.77%) changing the interpretation from P/LP to other categories, while only 64 of 1711 reclassified variants (inbound, 3.74%) were reclassified as P/LP. For those 64 variants that became P/LP, 34 variants were reclassified from conflicting interpretations, 28 variants were reclassified from VUS. Half of those were due to the addition of ENIGMA (QIMR Berghofer Medical Research Institute) as an expert panel submitter (according to ClinVar, interpretation from expert panel or practice guideline submitters overrules other interpretations). Only 2 B/LB variants were reclassified as P/LP. This resulted from re-interpretation of the same evidence - the ClinVar records submitted by OMIM (ClinVar # RCV000019494, and RCV000018360) were based on 1990s publications, which were misinterpreted as “benign” in the ClinVar August 2015 version. Overall, most of the re-classification in ClinVar feeds into “conflicting interpretation”, B/LB and VUS, and away from P/LP.

## DISCUSSION

It is known that the current knowledge on pathological variants is far from complete^29^. There have been several efforts to identify misclassification of variants including data sharing (e.g. ClinVar/ClinGen) and genetic analysis of large populations (e.g. identification of false positive interpretation for cardiomyopathy using 7855 cases^30^; re-evaluating previously identified casual variants for hypertrophic cardiomyopathy^31^). Such studies target and identify misclassified variants that are relatively common. Here, we not only identify conditions with relatively common variants but also conditions that have inflated genetic-risk (i.e. observed genetic risk is higher than population prevalence) when several rare variants are considered jointly.

This work revisits a well-understood relationship between minor allele frequency and disease prevalence. The pattern is one of an excess of individuals with genetic risk relative to the disease prevalence in the population. The inflation increases as the level of support for the genetic variants decreases in ClinVar. This means that a number of pathogenic variants have low penetrance or that those variants have an incorrect interpretation of pathogenicity. The present analyses strongly suggest that ClinVar includes significant amounts of misclassified variants, and supports the important role of ClinVar to increase transparency, contrast claims, and foster validation across submitters.

The ACMG-AMP standards and guidelines for variant interpretations recommends that variants with more than 5% allele frequency should be classified as benign variants (rule BA1)^3^. To lower this relatively lenient threshold, a more recent recommendation supports the use of disease-specific allele frequency (dAF) thresholds based on disease prevalence^1^. Application of the dAF threshold does correct inflation; few variants contribute to inflation for many of the conditions.

However, for some disorders inflation may be hard to pinpoint to low frequency alleles, and rare variants needed to be considered jointly. This problem is compounded by the very large number of rare variants that are being identified though genome and exome sequencing^10,32^. Here, rare variation risks being misconstrued as indicating functional relevance.

The present study also adds to the discussion on the importance of specifying review criteria, and improving the applicability of standardized criteria, such as those of the ACMG-AMP guidelines^1^. We include examples that illustrate some of the sources of misclassification and error and other issues in the reporting of pathogenic variants. Specifically, we observed cases of variants with low penetrance, incorrect mode of inheritance, conditions with unknown or older estimates of population prevalence, and variants that are incorrectly associated with disease.

We explored another strategy to understand the mechanisms of correction of the data currently deposited in ClinVar. Changes in classification in successive versions of ClinVar favors the general direction away from pathogenic/likely pathogenic towards VUS and benign/likely benign. However, the bulk of reclassified variants are reassigned to the “conflicting interpretation” category. This observation and our analysis of massive inflation in genetic risk for variants classified as having conflicting interpretation reinforce the notion that pathogenicity of these variants are questionable. More generally, the study supports the aim of reaching a definitive classification for this set of variants to avoid repeated re-assessment in the clinics.

In addition to classifying a variant as pathogenic or benign using ACMG-AMP guidelines, there is a need for a quantitative measure of risk (e.g. penetrance of the variant)^29^. Larger genetic studies integrating phenotype data to estimate variant risks using age/sex-specific incidence are needed. Although the concept of this work is anchored in the current practice of genetics, it serves to document the use of current large databases on the general population to better support the classification of variants. The observation of excessive numbers of individuals carrying genetic risk alleles both in well curated genes sets (ACMG-56) and in other resources (Orphanet) provides a useful benchmark for the improvement of variant annotation.

## Acknowledgments

All authors are either employees or consultants to Human Longevity, Inc.

